# Conspecific negative density dependence and why its study should not be abandoned

**DOI:** 10.1101/2020.05.11.089334

**Authors:** Joseph A. LaManna, Scott A. Mangan, Jonathan A. Myers

**Affiliations:** Department of Biological Sciences, Marquette University, PO Box 1881, Milwaukee, Wisconsin 53201 USA; Department of Biology & Tyson Research Center, Washington University in St. Louis, St. Louis, Missouri 63130 USA, &

**Keywords:** Community ecology, density dependence, dispersal, forest ecology, Janzen-Connell hypothesis, natural enemies, null models, spatial distribution, species interactions

## Abstract

Recent studies showing bias in the measurement of density dependence have the potential to sow confusion in the field of ecology. We provide clarity by elucidating key conceptual and statistical errors with the null-model approach used in Detto et al. (2019). We show that neither their null model nor a more biologically-appropriate null model reproduces differences in density-dependent recruitment between forests, indicating that the latitudinal gradient in negative density dependence is not an artefact of statistical bias. Finally, we suggest a path forward that combines observational comparisons of density dependence in multiple fitness components across localities with mechanistic and geographically-replicated experiments.

---

For over 50 years, ecological theory has hinged on the idea that self-suppression of populations caused by host-specific natural enemies or intraspecific competition contributes to patterns of species diversity at local, regional, and global scales (Janzen 1970, Connell 1971, Chesson 2000, Levine and HilleRisLambers 2009, Comita et al. 2010, Mangan et al. 2010, HilleRisLambers et al. 2012, LaManna et al. 2017, Usinowicz et al. 2017, Forrister et al. 2019). A key prediction of this hypothesis is that a species’ fitness should decline when or where it becomes common, thereby allowing rare species to persist in a community (i.e. negative frequency dependence). Observational studies conducted in temperate and tropical forests have tested this prediction by examining whether growth, survival, or recruitment of plants are reduced in areas of high conspecific density; a pattern known as conspecific negative density dependence, or CNDD (Harms et al. 2000, HilleRisLambers et al. 2002, Comita et al. 2010, LaManna et al. 2017, Johnson et al. 2012, Bagchi et al. 2014, Chen et al. 2019). For example, one study of large forest plots worldwide found that CNDD in tree recruitment – estimated as a decline in per-capita sapling recruitment with increasing conspecific adult density among localities within a forest – is stronger in tropical than temperate forests (LaManna et al. 2017). Such studies suggest that CNDD likely contributes to tree diversity around the globe. However, the utility of these approaches to address fundamental questions on the determinants of plant diversity has recently been called into question (Hülsmann and Hartig 2018, Detto et al. 2019). In particular, a recent paper by Detto et al. (2019) concluded that statistical bias alone likely accounts for most observed patterns of CNDD in plant communities, calling “into question the emerging paradigm that intraspecific competition has been demonstrated…to be generally stronger than interspecific competition” (pg. 1923). While we agree that careful consideration of bias is important in any observational analysis (Detto et al. 2019), the solution is not to abandon observational and phenomenological studies of CNDD. To help clear up confusion in the field, we clarify key conceptual and statistical problems with Detto et al. (2019), highlight the utility of evaluating multiple predictions of CNDD using different fitness components, and suggest a path forward for future studies of CNDD.

## Inference in observational studies: the importance of statistical replication of null models

Null models are important tools used to evaluate the importance of biotic interactions in ecological communities (Gotelli and Graves 1996). Yet the use of inappropriate null models can lead to flawed conclusions and confusion in the literature. For example, Detto et al. (2019) used a null model to claim that previous findings of stronger CNDD in tree recruitment in tropical than temperate forests (LaManna et al. 2017) were an artifact of statistical bias. However, the null-model analysis used by Detto et al. (2019) has major statistical and conceptual problems. The first problem is that their conclusions are based *on a single iteration of a relabeling null model*. By swapping “adult” and “sapling” labels among conspecifics while fixing their spatial distribution, they used relabeling to assess if observed differences in estimated CNDD between a tropical and temperate forest were due to biased estimates of adult and sapling densities, or spatial aggregation alone (Detto et al. 2019). However, a standard practice, and crucial step, in null-model analysis is to iterate the model hundreds to thousands of times to compare a distribution of simulated results to the observed result (Gotelli and Graves 1996). Using R-code and data from Detto et al. (2019), we calculated expected differences in median CNDD between the tropical and temperate forest for all species and rare species – the only difference was that we iterated their model 1,000 times. Contrary to conclusions in Detto et al. (2019), observed differences in CNDD between the tropical and temperate forest were larger than expected from the relabeling null model *once it was properly iterated* (Fig. 1).

**Figure 1.**
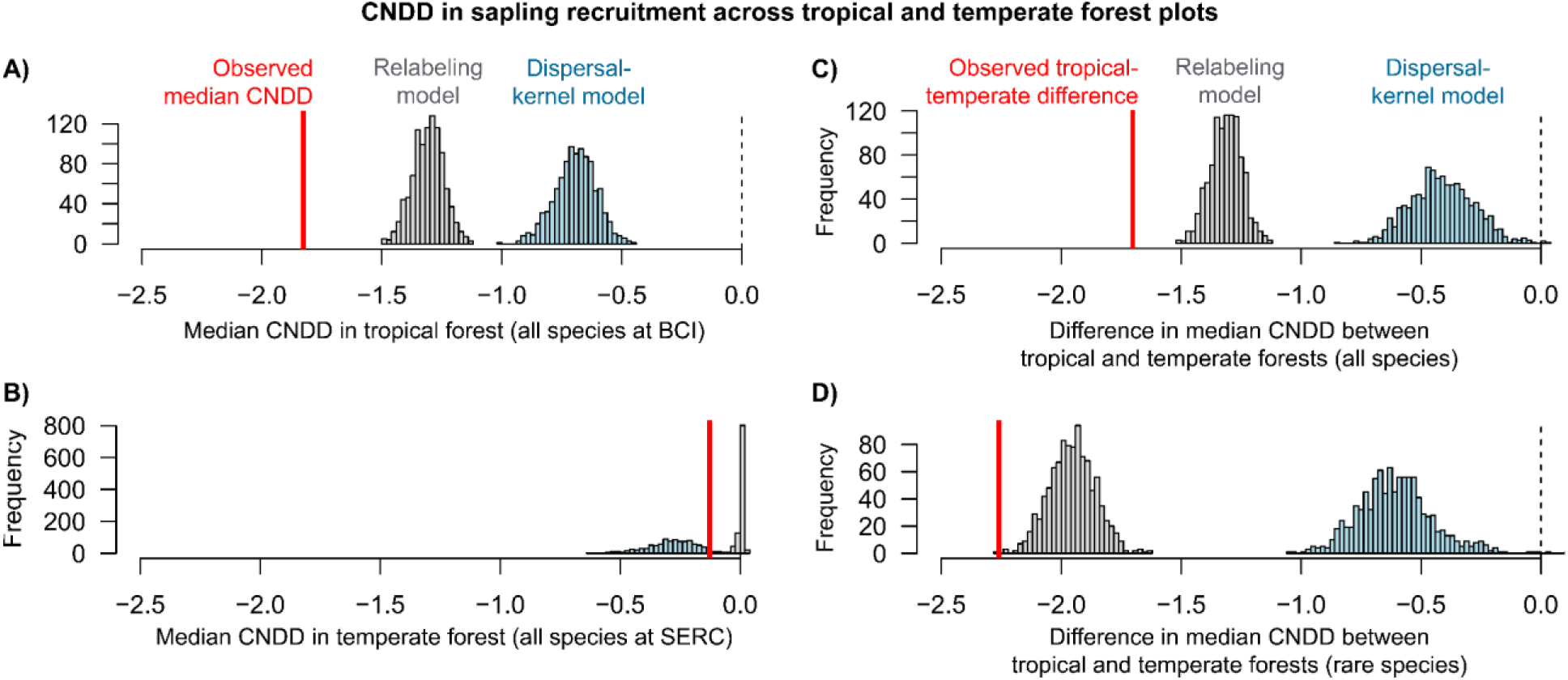
Re-analysis of data in Detto et al. (2019) from a tropical (BCI, Panama) and temperate (SERC, USA) forest. Observed results (red lines) are compared to null-expected results from 1,000 iterations of the relabeling model in Detto et al. (2019; gray histograms) and the dispersal-kernel model modified from LaManna et al. (2018) and described in the text (blue histograms; see acknowledgements for link to R code). (A) Observed median CNDD at BCI was greater than median CNDD expected from both models (*p* < 0.001). (B) Observed median CNDD at SERC was less than CNDD from the dispersal-kernel model (*p* = 0.036) but greater than CNDD from the relabeling model (*p* < 0.001). (C & D) Neither model reproduced the observed difference in median CNDD between tropical and temperate forests using (C) all species or (D) rare species with < 0.1 m^2^/ha basal area (*p* < 0.001). Ricker models and distance-weighted adult abundances were used in all analyses because they generated less bias in benchmark tests (LaManna et al. 2017, 2018b).

## Null models that fail to exclude hypothesized mechanisms can lead to flawed conclusions

The second major problem with the null-model analysis in Detto et al. (2019) is that it does not exclude the biological process of interest (i.e. the model is not density independent). As a result, their model induces false density dependence when there is none (Fig. 2). Because their relabeling model fixes the total number of individuals in a locality (e.g. quadrat of a forest plot), any area that is assigned a larger proportion of adults will necessarily be assigned a lower proportion of saplings (and vice versa), inducing a negative relationship between adult density and sapling recruitment. This failure to exclude density dependence causes relabeling to produce null patterns that are similar to observed estimates of CNDD (Fig. 1). In contrast, a more appropriate null model, such as the dispersal-kernel model used by LaManna et al. (2018a), does not induce false density dependence (Fig. 2). For this reason, dispersal-kernel models are recommended over relabeling models when evaluating any process that might affect spatial aggregation of recruits relative to adults (Wiegand and Moloney 2014). The dispersal-kernel model preserves adult locations to account for factors influencing adult aggregation (e.g. abiotic-habitat conditions), but excludes CNDD in recruitment by dispersing saplings according to empirically-supported estimates of seed-dispersal distances for each species (Thomson et al. 2011, Wiegand and Moloney 2014). This model also preserves and thereby accounts for differences in relative abundance and mean recruitment among species (LaManna et al. 2018a), and also accounts for potential influences of adult mortality by allowing a certain proportion of adults to die and be replaced by sapling recruits (see acknowledgements for link to R code). We applied the dispersal-kernel model to the same data used in Detto et al. (2019) and found even stronger support for the hypothesis that CNDD in recruitment is stronger in tropical than temperate forests (Fig. 1). This result is directly counter to results reported in Detto et al. (2019) and illustrates the pitfall of drawing conclusions from a biologically-inappropriate null model.

**Figure 2.**
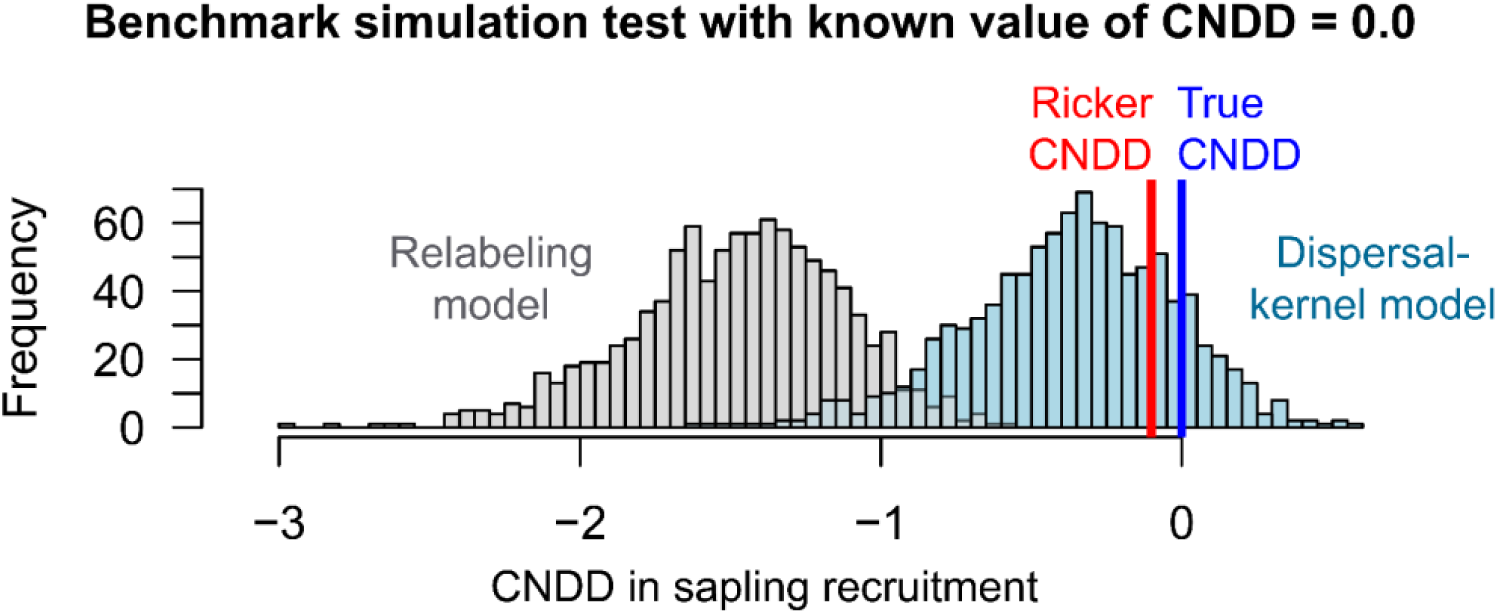
Benchmark simulation tests comparing 1,000 iterations of the relabeling model in Detto et al. (2019; gray histograms) and the dispersal-kernel model modified from LaManna et al. (2018) and described in the text (blue histograms; see acknowledgements for link to R code). Data is simulated for a hypothetical BCI species at relatively low abundance. Off-plot dispersal, adult mortality, and random error in sapling and adult abundances are also simulated here to reflect the influence of confounding processes. The known value of CNDD for these simulations is set to zero (blue line = truth), so an appropriate null model, like the dispersal-kernel model, should generate estimates that center on the observed CNDD estimate (red line = observed estimate from Ricker model). In contrast, the relabeling model generates strong, false density dependence when there is none – its null-value distribution does not even contain the observed CNDD estimate. This explains why the distribution of CNDD values from the relabeling model in analyses of real data are shifted to the left relative to the more appropriate dispersal-kernel model (Fig. 1). Ricker models and distance-weighted adult abundances were used in all analyses because they generated less bias in benchmark tests (LaManna et al. 2017, 2018b).

## Confounding biological factors can contribute to but not fully account for CNDD measures

All observational patterns in ecology are potentially subject to the influence of multiple forces, including spatial and temporal patterns predicted from CNDD. For example, spatial aggregation of conspecific individuals results from dispersal limitation, habitat selection, and any process that weakens CNDD in mortality or recruitment. Using two of the 24 forest plots analyzed in LaManna et al. (2017), Detto et al. (2019) found that species with more aggregated spatial distributions had weaker CNDD in recruitment as estimated in our study. Although this correlation supports the prediction that strong CNDD should decrease the aggregation of recruits around adults (Detto and Muller-Landau 2016, LaManna et al. 2017), Detto et al. (2019) conclude that these analyses are ambiguous because they do not account for other processes that influence aggregation of recruits. However, appropriate null models can account for the influence of other processes (Chave et al. 2002). In an earlier study that applied the dispersal-kernel model to all 24 forests plots (LaManna et al. 2018a), we found that the observed patterns in LaManna et al. (2017)—stronger CNDD in recruitment for rare tropical than temperate species and a latitudinal shift in the CNDD-abundance relationship—persist after accounting for species-specific estimates of dispersal and factors influencing adult aggregation (e.g., habitat selection). Additional analyses germane to key conclusions in Detto et al. (2019), including benchmark tests to explore influences of “error-prone proxies” and distance-weighted measures of adult abundance that minimize bias relative to quadrat-based measures, also show that findings in LaManna et al. (2017) persist after accounting for potential biases (LaManna et al. 2018a, 2018b). Interestingly, these finding suggest that latitudinal gradients in spatial aggregation cannot be understood without considering local biotic interactions underlying CNDD.

While dynamic measures of CNDD in survival or growth using multiple censuses are appealing because they are less influenced by dispersal, these measures are still susceptible to other confounding biological factors. For example, greater survival in areas of high conspecific density might not reflect weak density dependence but enhanced performance under favorable abiotic conditions. For these reasons, recent studies explicitly incorporate environmental parameters into models of CNDD and utilize appropriate null models to test the relative influence of hypothesized mechanisms (e.g. Zhu et al. 2015, Johnson et al. 2017). Thus, while Detto et al. (2019) suggest abandoning studies of CNDD based on spatial data, both temporal and spatial data suffer from the same problem – multiple non-mutually-exclusive biological processes can influence observed patterns. As in many other areas of science, the pursuit of a single measure that exists in isolation from any other influence is futile. No single “silver bullet” approach exists, but this does not mean that all observational studies in ecology, including the study of CNDD patterns, need to be abandoned. Instead, strong inference is still possible by utilizing appropriate null models and by integrating the effects of CNDD across multiple components of fitness.

## Different fitness components provide complementary insights into CNDD

CNDD enhances diversity via a combined influence on growth, survival, and recruitment that ultimately allows adults of one species to be replaced by adults of other species. However, studies that only examine CNDD in a single fitness component may completely miss important diversity-enhancing effects of CNDD on other fitness components or life stages (Bagchi et al. 2014, Zhu et al. 2015). Analyses of CNDD in growth, survival, and recruitment each have their own strengths and limitations (Zhu et al. 2015). An advantage of sapling-recruitment analyses is that they integrate effects of CNDD across multiple fitness components and early life-stages, including reproduction, seedling establishment, seedling survival and growth, and recruitment into the sapling age class (Zhu et al. 2015, LaManna et al. 2017). In contrast, growth and survival analyses offer several advantages over recruitment analyses (Comita et al. 2010, Detto et al. 2019), but do not evaluate CNDD effects on seed reproduction, germination, and recruitment into seedling and sapling age classes. Yet CNDD during early life stages may have disproportionate influences on diversity maintenance in later life stages (Comita et al. 2014, Green et al. 2014). Moreover, dynamic measures of CNDD usually examine changes in growth and survival over a short time relative to the long lifespans of most trees. Acknowledging these limitations is important because increased mortality at high conspecific densities alone will not enhance diversity if other species do not recruit in those areas (Janzen 1970, Connell 1971, Zhu et al. 2015). One promising approach is to separately measure CNDD in survival, growth and recruitment as a function of plant size and use integral projection models (IPMs; Merow et al. 2014, Levine et al. 2017) to determine the combined effect of CNDD on population growth rates. Studies that simultaneously examine CNDD across multiple life-stages and fitness components will be necessary to gain a more comprehensive understanding of CNDD’s effect on population and community dynamics.

## Towards a roadmap for future studies

Recent observational studies provide support for a latitudinal gradient in CNDD (Johnson et al. 2012, LaManna et al. 2017, 2018a, but see HilleRisLambers et al. 2002, Comita et al. 2014). Observational findings of a latitudinal gradient in CNDD are further supported by a growing number of replicated experimental studies of CNDD at different latitudes (Bagchi et al. 2014, Krishnadas et al. 2018, Jia et al. 2020). However, these studies have so far largely focused on only a few CNDD-generating mechanisms or on particular fitness components. Population and community dynamics will ultimately be influenced by differences in CNDD across localities and species regardless of the specific mechanisms generating that CNDD (Chesson 2000, Levine and HilleRisLambers 2009, HilleRisLambers et al. 2012). Compounding this problem is experimental evidence that several distinct mechanisms can generate CNDD, including interactions with host-specific pathogens or insects, and strong intraspecific competition (Mangan et al. 2010, McCarthy-Neumann and Kobe 2010, Bagchi et al. 2014, Jia et al. 2020). The relative importance of these mechanisms also differs among species (Jia et al. 2020), meaning that even a carefully-designed study examining one or a few mechanisms, while valuable, would nonetheless fail to detect CNDD caused by other mechanisms. Large-scale observational studies are therefore a key first step to elucidate whether the demographic signatures of CNDD exist in nature, whether they vary among localities and species, and inform the design of experiments. Indeed, such pattern-based studies are an intermediate link between specific mechanisms generating CNDD, which are myriad, and fundamental community properties such as species diversity and relative abundance.

A productive way forward will be to integrate mechanism-based and pattern-based approaches. Specifically, we propose a two-pronged approach: 1) first test for phenomenological patterns predicted by CNDD at different scales of biological organization (individual growth and mortality, population vital rates, community dynamics), and then 2) explore mechanisms underlying those patterns using experimental studies of competition and natural enemies that have the potential to be geographically replicated (e.g. Forrister et al. 2019, Jia et al. 2020). For example, phenomenological studies of CNDD in tropical forests have highlighted the importance of interacting species (Comita et al. 2010, Usinowicz et al. 2017, Johnson et al. 2017), and recent studies of mechanisms showed that several agents, including fungal pathogens and insects, are likely to be a driving force behind these patterns (Mangan et al. 2010, Bagchi et al. 2014, Gripenberg et al. 2019, Forrister et al. 2019). Future observational and experimental tests of multiple predictions of CNDD across tropical and temperate forests will provide key insights into how local-species interactions influence large-scale diversity gradients.

## Acknowledgements

We thank Stefan Schnitzer, Liza Comita, an anonymous reviewer, and members of the Schnitzer and LaManna labs for helpful comments. We also thank the many scientists responsible for collecting Forest Global Earth Observatory (ForestGEO) data, and the National Science Foundation for supporting our research (DEB-2024903 to J.A.L.; DEB-1557094 to J.A.M.; DEB-1257989 to S.A.M.). R-code and links to data to reproduce our analyses can be found at: https://github.com/jalamanna/Comment-on-Bias-in-the-detection-of-negative-density-dependence.

## Notes

https://github.com/jalamanna/Comment-on-Bias-in-the-detection-of-negative-density-dependence

### Competing Interest Statement

The authors have declared no competing interest.

## References

Bagchi, R., R. E. Gallery, S. Gripenberg, S. J. Gurr, L. Narayan, C. E. Addis, R. P. Freckleton, and O. T. Lewis. 2014. Pathogens and insect herbivores drive rainforest plant diversity and composition. Nature 506:85–88.

Chave, J., H. C. Muller-Landau, and S. A. Levin. 2002. Comparing classical community models: theoretical consequences for patterns of diversity. The American Naturalist 159:1–23.

Chen, L., N. G. Swenson, N. Ji, X. Mi, H. Ren, L. Guo, and K. Ma. 2019. Differential soil fungus accumulation and density dependence of trees in a subtropical forest. Science 366:124–128.

Chesson, P. 2000. Mechanisms of maintenance of species diversity. Annual Review of Ecology and Systematics 31:343–366.

Comita, L. S., H. C. Muller-Landau, S. Aguilar, and S. P. Hubbell. 2010. Asymmetric density dependence shapes species abundances in a tropical tree community. Science (New York, N.Y.) 329:330–332.

Comita, L. S., S. A. Queenborough, S. J. Murphy, J. L. Eck, K. Xu, M. Krishnadas, N. Beckman, and Y. Zhu. 2014. Testing predictions of the Janzen–Connell hypothesis: a meta-analysis of experimental evidence for distance-and density-dependent seed and seedling survival. Journal of Ecology 102:845–856.

Connell, J. H. 1971. On the role of natural enemies in preventing competitive exclusion in some marine animals and in rain forest trees. Pages 298–312 *in* P. J. den Boer and G. R. Gradwell, editors. Dynamics of populations. Centre for Agricultural Publishing and Documentation, Wageningen, The Netherlands.

Detto, M., and H. C. Muller-Landau. 2016. Rates of formation and dissipation of clumping reveal lagged responses in tropical tree populations. Ecology 97:1170–1181.

Detto, M., M. D. Visser, S. J. Wright, and S. W. Pacala. 2019. Bias in the detection of negative density dependence in plant communities. Ecology Letters 22:1923–1939.

Forrister, D. L., M.-J. Endara, G. C. Younkin, P. D. Coley, and T. A. Kursar. 2019. Herbivores as drivers of negative density dependence in tropical forest saplings. Science 363:1213–1216.

Gotelli, N. J., and G. R. Graves. 1996. Null models in ecology. Smithsonian Institution, Washington, D.C., USA.

Green, P. T., K. E. Harms, and J. H. Connell. 2014. Nonrandom, diversifying processes are disproportionately strong in the smallest size classes of a tropical forest. Proceedings of the National Academy of Sciences of the United States of America 111:18649–18654.

Gripenberg, S., Y. Basset, O. T. Lewis, J. C. D. Terry, S. J. Wright, I. Simón, D. C. Fernández, M. Cedeño-Sanchez, M. Rivera, H. Barrios, J. W. Brown, O. Calderón, A. I. Cognato, J. Kim, S. E. Miller, G. E. Morse, S. Pinzón-Navarro, D. L. J. Quicke, R. K. Robbins, J.-P. Salminen, and E. Vesterinen. 2019. A highly resolved food web for insect seed predators in a species-rich tropical forest. Ecology Letters 22:1638–1649.

Harms, K. E., S. J. Wright, O. Calderón, A. Hernández, and E. A. Herre. 2000. Pervasive density-dependent recruitment enhances seedling diversity in a tropical forest. Nature 404:493–495.

HilleRisLambers, J., P. B. Adler, W. S. Harpole, J. M. Levine, and M. M. Mayfield. 2012. Rethinking community assembly through the lens of coexistence theory. Annual Review of Ecology, Evolution, and Systematics 43:227–248.

HilleRisLambers, J., J. S. Clark, and B. Beckage. 2002. Density-dependent mortality and the latitudinal gradient in species diversity. Nature 417:732–735.

Hülsmann, L., and F. Hartig. 2018. Comment on “Plant diversity increases with the strength of negative density dependence at the global scale.” Science 360:eaar2435.

Janzen, D. H. 1970. Herbivores and the number of tree species in tropical forests. American naturalist 104:501–528.

Jia, S., X. Wang, Z. Yuan, F. Lin, J. Ye, G. Lin, Z. Hao, and R. Bagchi. 2020. Tree species traits affect which natural enemies drive the Janzen-Connell effect in a temperate forest. Nature Communications 11:1–9.

Johnson, D. J., W. T. Beaulieu, J. D. Bever, and K. Clay. 2012. Conspecific negative density dependence and forest diversity. Science (New York, N.Y.) 336:904–907.

Johnson, D. J., R. Condit, S. P. Hubbell, and L. S. Comita. 2017. Abiotic niche partitioning and negative density dependence drive tree seedling survival in a tropical forest. Proceedings of the Royal Society B: Biological Sciences 284:20172210.

Krishnadas, M., R. Bagchi, S. Sridhara, and L. S. Comita. 2018. Weaker plant-enemy interactions decrease tree seedling diversity with edge-effects in a fragmented tropical forest. Nature Communications 9:1–7.

LaManna, J. A., S. A. Mangan, A. Alonso, et al. 2017. Plant diversity increases with the strength of negative density dependence at the global scale. Science (New York, N.Y.) 356:1389–1392.

LaManna, J. A., S. A. Mangan, A. Alonso, et al. 2018a. Response to Comment on “Plant diversity increases with the strength of negative density dependence at the global scale.” Science 360:eaar3824.

LaManna, J. A., S. A. Mangan, A. Alonso, et al. 2018b. Response to Comment on “Plant diversity increases with the strength of negative density dependence at the global scale.” Science 360:eaar5245.

Levine, J. M., J. Bascompte, P. B. Adler, and S. Allesina. 2017. Beyond pairwise mechanisms of species coexistence in complex communities. Nature 546:56–64.

Levine, J. M., and J. HilleRisLambers. 2009. The importance of niches for the maintenance of species diversity. Nature 461:254–257.

Mangan, S. A., S. A. Schnitzer, E. A. Herre, K. M. Mack, M. C. Valencia, E. I. Sanchez, and J. D. Bever. 2010. Negative plant-soil feedback predicts tree-species relative abundance in a tropical forest. Nature 466:752–755.

McCarthy-Neumann, S., and R. K. Kobe. 2010. Conspecific and heterospecific plant–soil feedbacks influence survivorship and growth of temperate tree seedlings. Journal of Ecology 98:408–418.

Merow, C., J. P. Dahlgren, C. J. E. Metcalf, D. Z. Childs, M. E. Evans, E. Jongejans, S. Record, M. Rees, R. Salguero-Gómez, and S. M. McMahon. 2014. Advancing population ecology with integral projection models: a practical guide. Methods in Ecology and Evolution 5:99–110.

Thomson, F. J., A. T. Moles, T. D. Auld, and R. T. Kingsford. 2011. Seed dispersal distance is more strongly correlated with plant height than with seed mass. Journal of Ecology 99:1299–1307.

Usinowicz, J., C.-H. Chang-Yang, Y.-Y. Chen, J. S. Clark, C. Fletcher, N. C. Garwood, Z. Hao, J. Johnstone, Y. Lin, M. R. Metz, T. Masaki, T. Nakashizuka, I.-F. Sun, R. Valencia, Y. Wang, J. K. Zimmerman, A. R. Ives, and S. J. Wright. 2017. Temporal coexistence mechanisms contribute to the latitudinal gradient in forest diversity. Nature 550:105–108.

Wiegand, T., and K. A. Moloney. 2014. Handbook of spatial point-pattern analysis in ecology. CRC Press, Boca Raton, Florida, USA.

Zhu, K., C. W. Woodall, J. V. Monteiro, and J. S. Clark. 2015. Prevalence and strength of density-dependent tree recruitment. Ecology 96:2319–2327.

